# Directed Brain Connectomics Revealed by Bicommunity Structure

**DOI:** 10.64898/2026.02.25.707616

**Authors:** Alexandre Cionca, Chun Hei Michael Chan, Francesca Saviola, Maciej Jedynak, Yasser Alemán-Gómez, Saina Asadi, Arthur Spencer, Olivier David, Ileana Jelescu, Patric Hagmann, Maria Giulia Preti, Dimitri Van De Ville

## Abstract

The brain is a complex, interconnected system in which structural wiring underpins the flow of information between functional units. After two decades of advances in connectomics, the asymmetry of edge structures in large-scale brain networks remains largely unexplored. Here, we build a directed connectome combining structure and function, and dissect its organization into bicommunities where sending and associated receiving sets of nodes, through clustering of directed-edge information, are fully acknowledged. Our findings reveal a primary directional axis from sensory to association cortices, indicative of bottom-up information flow. Validation with invasive electrophysiological measures demonstrates that these bicommunities capture an intermediate organizational scale at which connection asymmetry aligns across modalities. Finally, sets of distinct known anatomical fiber bundles and their functional mapping are recovered as specific bicommunities. This work identifies asymmetric pathways onto which neural computation can be expressed, and establishes directed connectomics as a novel framework for understanding brain organization.

## Introduction

Human brain connectomics refers to the study of the tangled maps of large-scale neural connections; how they are organized and how they support brain function and ultimately cognition and behavior (1,2). The wiring diagram reflected in the structural connectome can be complemented by a functional one that captures statistical interdependences of neural activity in pairs of regions. Driven by advances in network science and computational methods, the exploration of brain networks using graph modeling led to insightful findings about its networked hierarchy into hubs, modules of densely interconnected regions, and principal axes or gradients (e.g., the sensori-association axis from unimodal to transmodal regions), that can be used for phenotyping of groups and individuals (3–7). One recurrent observation is that connectomes are organized into distinct yet interacting communities in which regions are densely interconnected in terms of functional or structural connectivity (4,5). Comparable modular structures have been observed in other species, such as the *Caenorhabditis elegans*, *Drosophila* fruit fly or macaque, suggesting that modularity is evolutionary preserved across nervous systems (8–11).

The current mesoscale description of the human connectome captures global aspects of brain architecture, but remains oblivious to the degree of directionality of the axonal fibers giving rise to asymmetry and hierarchy in information flow within the brain (12–14). Non-invasive imaging techniques cannot resolve directionality of structural connections; i.e., diffusion-weighted MRI and tractography-based reconstructions of white matter bundles only render undirected connectomes (15). Nevertheless, computational and biophysical models provide estimates of directed connectivity from temporal features of brain activity (16–19), structural topology (20,21), or both (22,23). First, using functional data, approaches such as dynamical causal modeling (DCM) or Granger causality infer the direction of influence in time series by exploiting temporal precedence, thereby identifying sources that drive the observed dynamics. While limitations of such models, such as scalability, have been mitigated, for instance, through regression DCM (rDCM) that incorporates structural priors (23), effective connectivity (EC) face uncertain neurobiological validity, alongside the requirement for long time series (23,24). Second, with the help of network communication models, the topological wiring of the human brain can provide evidence of asymmetric routing from undirected connectomes (20,21). Whether from a hierarchical or polysynaptic (multi-hop) routing perspective, these frameworks demonstrate that directionality is intrinsic to the architecture of brain graphs and that connection asymmetry mirrors the canonical sensory-associative axis (6,20). Third, structure-informed models of directed connectivity bridge structural and temporal approaches by leveraging white-matter anatomy to, for example, restrict the parameter space in rDCM (23,25) or penalize structurally implausible connections in time-varying multivariate autoregressive (tvMVAR) models (26,27). However, structure-aware approaches are usually indirectly validated against effective or functional connectivity and, especially for MVAR, may not be accurate with low temporal resolution, such as resting-state fMRI (22,26). Mapping whole-brain directed connectivity into large-scale networks thus remains a critical challenge, constrained by methodological limitations and that lack consensus and data to validate directed connectomes.

In fact, directed human connectomes scarcely include validation using measurements of neuronal propagation, whereas studies in animal models have bridged this validation gap through the use of invasive methods such as optogenetics, retrograde tracers or intracranial recordings (8,28–30). Yet, there is a recent emergence of human invasive datasets that include, for example, intracranial/stereo electroencephalography (iEEG/sEEG) in patients with epilepsy (12,14,31,32). They provide sparse but direct measurements of asymmetrical neuronal conductions, which, at a population level, approximate large-scale directed communication across neuronal ensembles. By combining such datasets in a common framework that integrates multimodal and multiscale organizations, one could hope to validate the directional hierarchy in human connectomes (22,33).

Traditionally, network neuroscience has been centered on modeling the brain as a collection of functionally similar nodes, which, in cases, can be informed with directionality (34,35). Indeed, the asymmetry of connections, as usually captured by the nodal in- and out-degree, is a defining feature of brain communication and reflects macroscale functional specialization, which is ultimately tied to its cytoarchitecture (14,36–38). However, such nodal aggregates may blur important aspects of the underlying dynamics, as neural interactions are more likely to have subtle directional biases than pure unidirectional signaling. It appears a paradigm shift is underway: from exploring how to group regions into communities to understanding which are the driving pathways, the focus is essentially moving from nodal to edge, or higher-order representations (39–42). Dissection of the human connectome from an edge-centric perspective shows that the brain organization extends beyond conventional assortative communities (i.e., densely connected nodes) to include disassortative (bipartite graph behavior) and core-periphery (mixture of both extremes) communities (39,43). Circling back to directed graphs, we stress the need to consider non-assortative communication as it inherently encodes asymmetric flow from distinct source to target nodes (44). In short, the shift from nodes to edges, and from undirected to directed graphs, goes beyond a pure methodological extension and raises a conceptual reorientation: the computational architecture of the brain is built on asymmetric pathways, where the functional role of each region is then paralleled by how information flows through them.

The contribution of this work is twofold: first, we derive a human whole-brain directed structural connectome by combining diffusion MRI for the undirected structural backbone and rDCM of fMRI to infer directionality (35,45); second, we leverage a novel framework for detecting directed graph communities, named *bicommunities* (42), and explore the hierarchical organization of the directed brain network. In essence, bicommunities reinterpret graph communities as a set of coordinated, directed pathways instead of densely interconnected nodes. Through the detection of clusters of graph edges, we identify coherent white matter pathways with preferred directionality and with corresponding bundles that are mainly along association fibers. Furthermore, connections between the hemispheres or along projection fibers remain bidirectional. We validate the observed directionality by incorporating invasive cortical recordings of electrical propagation between pairs of brain regions (31). In detail, we found high similarities between edge direction asymmetry in the directed connectome and in conduction delays when seen through the lens of the bicommunities. Our edge-centric perspective distinguishes sending and receiving patterns of assortative and core-periphery communication from purely disassortative bicommunities that, for example, connect transmodal areas along the arcuate and superior lateral fasciculus. Finally, in an effort to bridge brain structure and function, we summarize how groups of known anatomical bundles may connect resting-state networks. Our work unifies directed information at multiple temporal scales through a common structural scaffold, the bicommunities, and thus places directed connectomics as a key methodological avenue to better comprehend the complexity of neuronal wiring.

## Results

### Hierarchical Clustering of the Directed Connectome

We start by building a directed connectome defined as the structural wiring of white matter fibers (45), a binary graph, informed with directionality (Fig. 1. A). Directionality is defined as the edge-wise asymmetry observed in a whole-brain estimate of effective connectivity (EC) through the regression dynamical causal modelling (rDCM) framework applied to resting-state functional MRI (23,35). In this context, edge asymmetry, refers to the imbalance between the incoming and outgoing edge weight of the same connection (defined as the normalized ratio of outgoing weight over the sum of bidirectional weights). The resulting connectome is made of 129 nodes and 7906 edges, where 25% of connections have a directionality higher than 0.54 and 1.26% are strictly directed (edge asymmetry = 1, Fig. S1). We then explore the directed organization of the brain through our recent framework based on bimodularity for the detection of directed communities (42). Briefly, this approach identifies groups of edges, termed bicommunities, that capture sets of corresponding sending and receiving nodes by features in an embedding space. We ensure stability and robustness of the clustering scheme through hierarchical clustering applied to the consensus matrix (Fig. 1. B-C). This matrix captures how often two edges are paired in a similar cluster for multiple instances of K-means and over a wide range of number of clusters *K*. We observe stable bicommunities for settings that gradually increase from coarse (*K*=11) to fine (*K*=75) separation of the graph edges and show three local maxima (Fig. 1. C) of branching distance that are well within the range of high consensus stability (Fig. S2). Regardless of the number of clusters, the bicommunities capture directed connectivity between and within specific brain lobes and thus summarize thousands of edges into dozens of principal pathways (Fig. 1. D-F, S3). Expressed with proportions, compressing 7906 connections into *K*=35 bicommunities (most stable cluster configuration) corresponds to a size reduction factor of 225. Bicommunities distinguish the commissural from the association fibers as well as known anatomical bundles, such as the arcuate and superior longitudinal fasciculi, which are captured within distinct clusters of edges with different levels of granularity (Fig. 1. G-I, S4). We observe that most bicommunities have a counterpart, a conjugate, such that the edges in the bicommunity and its conjugate are oriented in opposite directions. In these conjugate pairs, the sending nodes of one can be similar to the receiving nodes of the other, thus describing both directions of a same pathway in which edge weights may not be similar, thus reflecting a preferred direction. It is also noticeable that unilateral bicommunities (limited to one hemisphere) tend to have a corresponding bicommunity on the other hemisphere, reflecting well-known brain symmetric organization between homologous regions.

**Figure 1.**
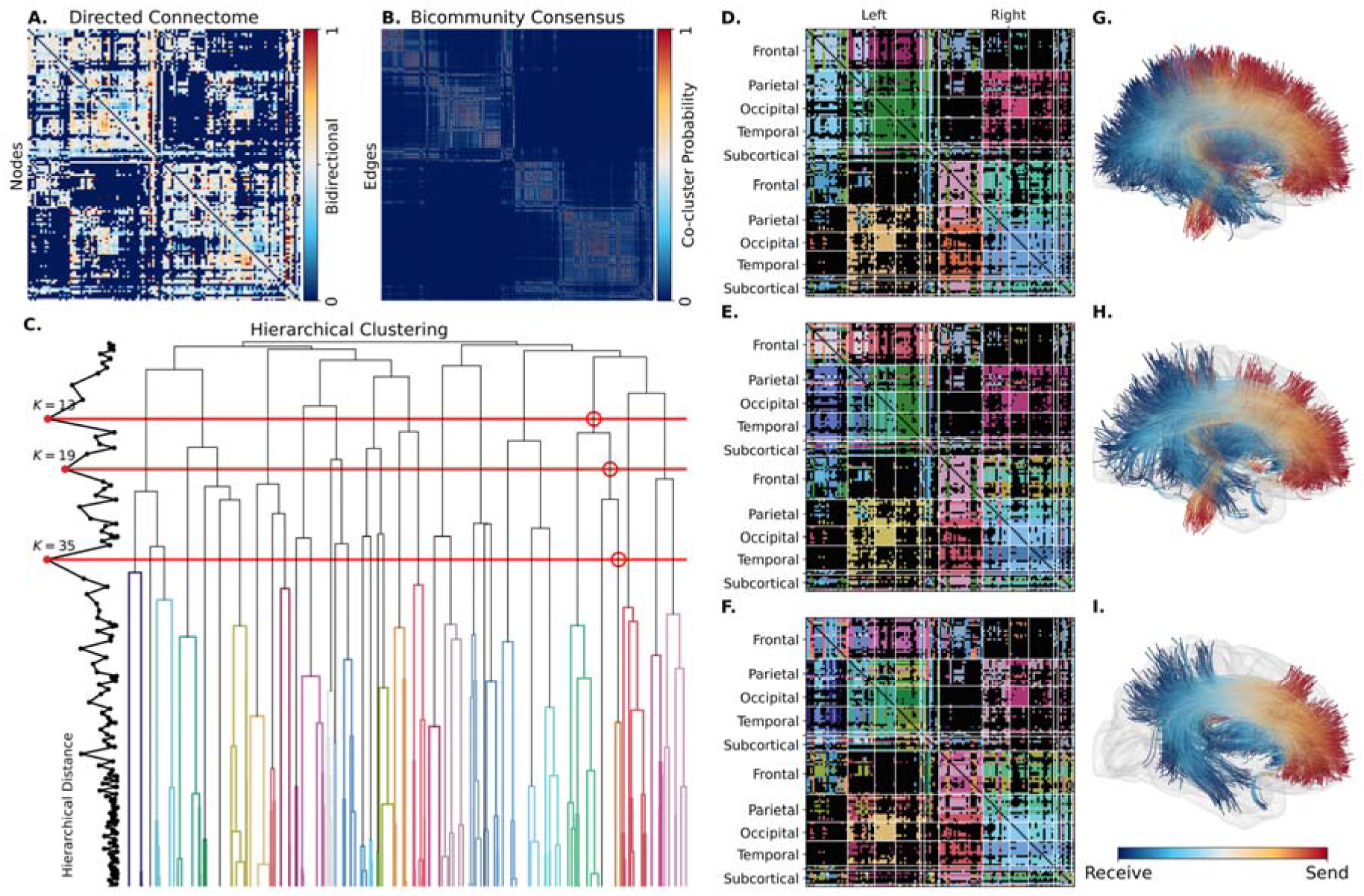
Hierarchical Clustering of the Directed Connectome. **A)** Directed structural connectivity matrix. **B)** Consensus clustering matrix from the detection of bicommunities. Each entry represents the K-means occurrences in which two edges were found in the same cluster. **C)** Branching distance and dendrogram of the consensus clustering matrix with the 3 highest local maxima (*K*=13, *K*=19, *K*=35) that indicate stable clustering. Red horizontal lines indicate the cutting distances in the hierarchical tree and red circles highlight the bicommunities selected in **D-F**. Colored lines correspond to edge clusters for *K*=35 (**F**, **I**). **D-F)** Colored connectivity matrices showing the edge clusters where each color denotes a bicommunity for each of the 3 local maxima (**D**: *K*=13, **E**: *K*=19, **F**: *K*=35). **G-I)** Streamline representation of selected bicommunities (ordered as in **D-F**) from the same hierarchical branch as indicated by the red circles in **C**. Centroids of white matter streamlines for each edge (5 centroids per edge) that composes the bicommunity are aggregated and colored to indicate the sending and receiving end of each streamline bundle.

### Preferred Directionality of White-Matter Pathways

Directionality is intrinsic to the detection of bicommunities as it groups edges that identify sets of sending and receiving nodes that typically do not overlap. It is however not apparent if edges within a bicommunity are bidirectional or if the associated white matter pathway has a preferred direction. As such, we estimate the presence of such a preferred direction by evaluating the degree of edge asymmetry within a bicommunity. The statistical significance of edge asymmetry is assessed through comparison with a null distribution computed by randomly swapping (N_Permutations_=10000) the directional weight of each edge, under the null hypothesis that the directional information is spurious. Fixing *K*=35, we identify 14 bicommunities (40%) with edge weight asymmetry that significantly differ from 0.5 (bidirectional edges) with p-values < 0.05 after Bonferroni-correction for multiple tests (N_Tests_=*K*=35, Fig. 2. B). Bicommunities with preferred directions are mostly found along association fibers and connecting regions of a same hemisphere, whereas commissural fibers appear bidirectional (Fig. 2. A). These preferred directions appear to follow a main axis of directed communication pathways from temporal/somatosensory to occipital networks and back from frontal to occipito-temporal areas (Fig. 2. C). On the other hand, bidirectional pathways mainly group commissural edges and capture subparts of the corpus callosum (Fig. 2. D) hinting that there might not be significant directionality in interhemispheric connections. We find similar proportions of edge clusters with preferred direction when varying the number of clusters *K* (Fig. S2) and the associated axes of directed communication are coherent (Fig. S5, S6).

**Figure 2.**
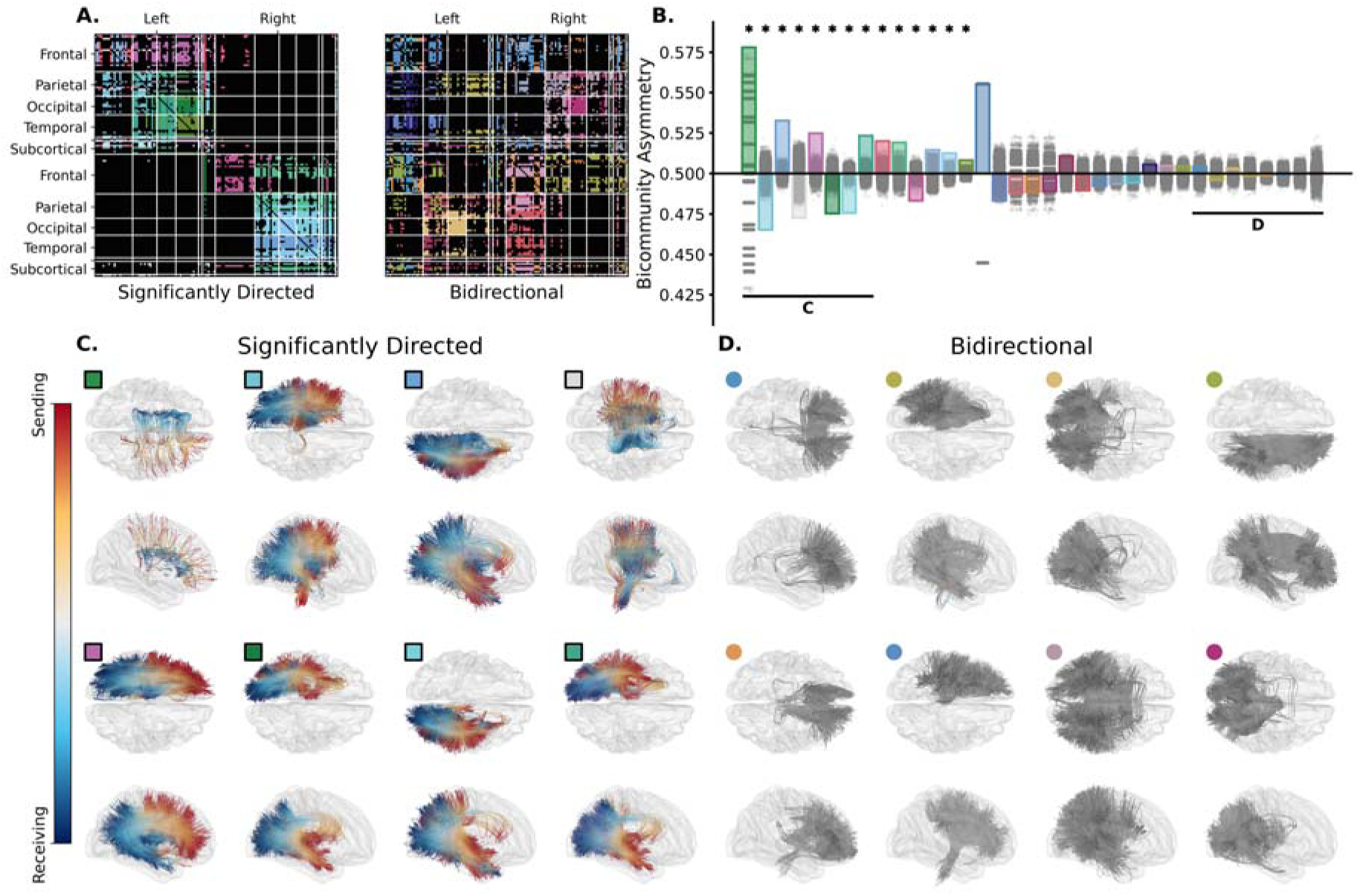
Directionality of White-Matter Pathways. **A)** Edge cluster matrix showing the cluster assignment of edges to individual bicommunities separated in significantly directed (left) or non-significant (right) edge asymmetry. Graph nodes are ordered by brain lobes (rows) and hemisphere (columns). **B)** Bar plots show the median edge asymmetry within each of the *K*=35 bicommunities with colors to indicate edge to cluster assignment as in **A**. Gray scatter points show the null distribution of asymmetry after reshuffling edge weight directions, and dark asterisks show significantly directed bicommunities (p-value < 0.05 after Bonferroni correction). **C-D)** White matter streamline centroids (5 per edge) are shown for the 8 bicommunities with the highest (**C**) and lowest (**D**) median directionality (as highlighted with two horizontal lines in **B**) in the transverse (first and third rows) and sagittal (second and fourth rows). Colored markers show the bicommunity correspondence with **A** and **B** where squared and circle markers indicate significant and non-significant edge asymmetry respectively. Streamlines are colored based on which end is sending (red) and receiving (blue), or fully in gray for bidirectional bicommunities. Streamline directions have been adapted to reflect the preferred direction (flipped if edge asymmetry is < 0.5).

### Invasive Validation of Connectome Directionality

As a validation of the directed connectome and to ensure that the observed directionality is meaningful, we propose a comparison with intracranial electrophysiological measurements, part of the F-Tract dataset (31). In particular, as measured through stereo-encephalography (sEEG), cortico-cortical evoked potentials (CCEPs) capture the delayed response to a direct electrical stimulation between electrodes at different locations, and can estimate, among other features, the delay of axonal propagation. We consider different temporal metrics such as the time between stimulation and: 1) the first local minimum before response (latency start); 2) the passing of a statistical threshold (onset delay); 3) and the first peak of the response (peak delay). The amplitude of response (z score with respect to the baseline) is also included as part of the validation features. Therefore, these in-vivo measurements of brain communication provide a “ground truth” for edge directionality in which delays and response amplitude capture the efficiency and strength of directed connections, respectively. F-Tract metrics are then summarized for each region pair in the connectome atlas, resulting in directed connectivity matrices in which edge-wise asymmetry can be estimated as for the effective connectivity. Asymmetry in delays thus reflects conduction speed (lower is faster) and asymmetry in the response amplitude quantifies the directional imbalance in the amount of transmitted information (higher means more signal). First, we naively compare the edge asymmetry between F-Tract and our connectome at the level of individual connection which reveals poor agreement (Spearman’s |ρ| < 0.2) between the two modalities (Fig. 3. A). Second, we assess if the grouping of edges given by the bicommunities capture directional information that can be reflected in CCEPs. For each of the F-Tract features, we compute the median asymmetry connections within each bicommunity and compare this aggregated directionality with the one derived from our connectome. We observe a moderate agreement (Spearman’s |ρ| > 0.4) between the two modalities and, for all F-Tract measurements, there is a higher correlation when the comparison is made at the level of bicommunities than in the single-edge approach (Fig. 3. B). We note that the observed correlations between F-Tract and the connectome asymmetry are negative for temporal delays and positive for response amplitude, which is coherent with long delays being associated with weaker connectivity. Furthermore, and to reject the hypothesis that high similarity may be found when considering any group of edges, we repeat the computation of Spearman correlation in bicommunity settings in which the cluster assignment of graph edges has been reshuffled (N_Permutations_=4999). This builds a null distribution of correlations between edge asymmetry in the connectome and in F-Tract features for randomized bicommunity structure. We observe that agreements between CCEPs and the connectome directionality are higher than in random settings for specific numbers of bicommunities (Fig. 3. B). Coarser clustering of the graph edges (*K*=19) show significantly higher correlations in more F-Tract features than finer ones (*K*=35). Yet, all three stable configurations of bicommunity structure capture substantial parts of the directed information in F-Tract, for both delays in electrical propagation and response amplitude.

**Figure 3.**
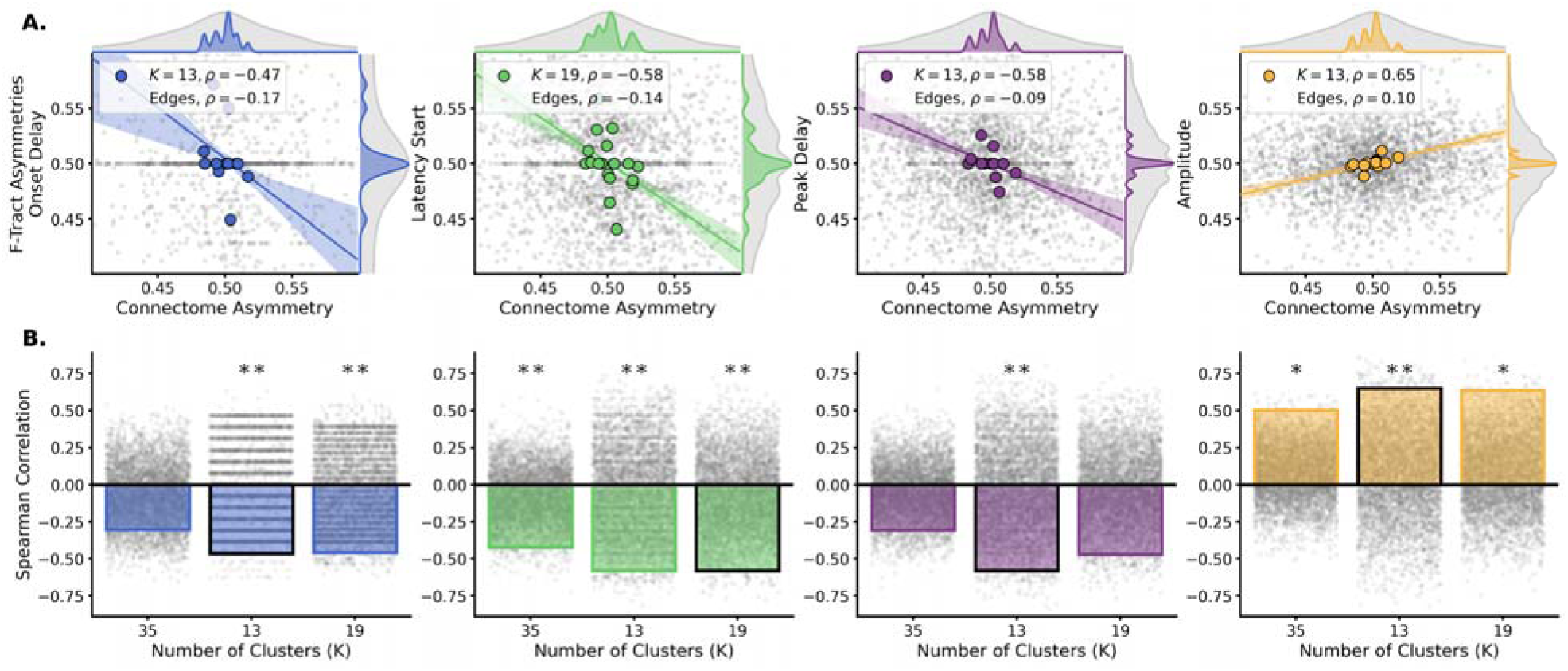
Agreement Between Edge Asymmetry across Modalities. **A)** Agreement, as captured by scatter-plots and Spearman correlation, between the edge asymmetry in the connectome (horizontal) and selected F-Tract features (vertical). Scatter and correlation are shown for comparison at the level of the graph edges (gray dots, “Edges” legend) and of the bicommunities (colored dots, legend with *K* being the number of edge clusters). A linear regression line is shown in color along with the minimum-maximum bound (filled area) of regression lines when removing individual bicommunities (leave-one-out). The outer curves show the distribution of edge asymmetry for F-Tract connectivity (right) and for our connectome (top) when considering graph edges (gray) or bicommunities (colored). **B)** Bars show the Spearman correlation between the median asymmetry of the connectome and F-Tract feature at the level of the bicommunities for the three local maxima of hierarchical clustering stability (*K*=35, *K*=13, *K*=19). Scattered dots show the null distribution of correlation with shuffled edge cluster assignment (4999 permutations). The bar highlighted with a dark stroke indicates the configuration shown in the scatter in **A**. * p < 0.05 uncorrected, ** p < 0.05 FDR corrected for multiple tests.

### Situating Bicommunities on the Assortative to Disassortative Spectrum

To investigate the network properties of bicommunities, we characterize directed graph communities by considering their assortativity and tendencies either to send or receive. Assortative communities are densely connected nodes (i.e., conventional communities) and disassortative communities indicate dense bipartite graph behavior (i.e., strong connectivity between two disjoint sets of nodes) (43). Furthermore, assortativity is a spectrum and there are more nuanced patterns of communications such as core-periphery communities. These are similar to assortative connections between nodes, but with the inclusion of disassortative edges going to (or coming from) peripheral nodes. For the case of *K*=35 bicommunities, we observe that most of the clusters of edges with preferred direction tend to be on the assortative end of the spectrum (Fig. 4). There is, however, a distinction between these core-periphery patterns that are either sending or receiving dominant. For example, projection from primary sensory to frontal areas or from temporal to occipital lobe exhibit a sending core-periphery type of communication in which there are more sending regions than receiving ones (Fig. 4, top right box). In the other hemisphere, the opposite pattern is found in similar connections from somatosensory to frontal area but where the proportion of sending and receiving nodes has been swapped (Fig. 4, bottom right box). Considering the disassortative half of the axis (Fig. 4, left boxes), we observe a communication pathway from the temporal and parietal area that appear to focus information in the occipital lobe and which appears in both hemispheres. Furthermore, purely disassortative bicommunities mainly capture communication along commissural fibers (through the corpus callosum in our case) without preferred direction. Nevertheless, we observe assortative inter-hemispheric connections within the superior frontal gyri and without directionality (Fig. 4, top right box), hinting that homotopic commissural fibers can, in cases, form densely connected communities.

**Figure 4.**
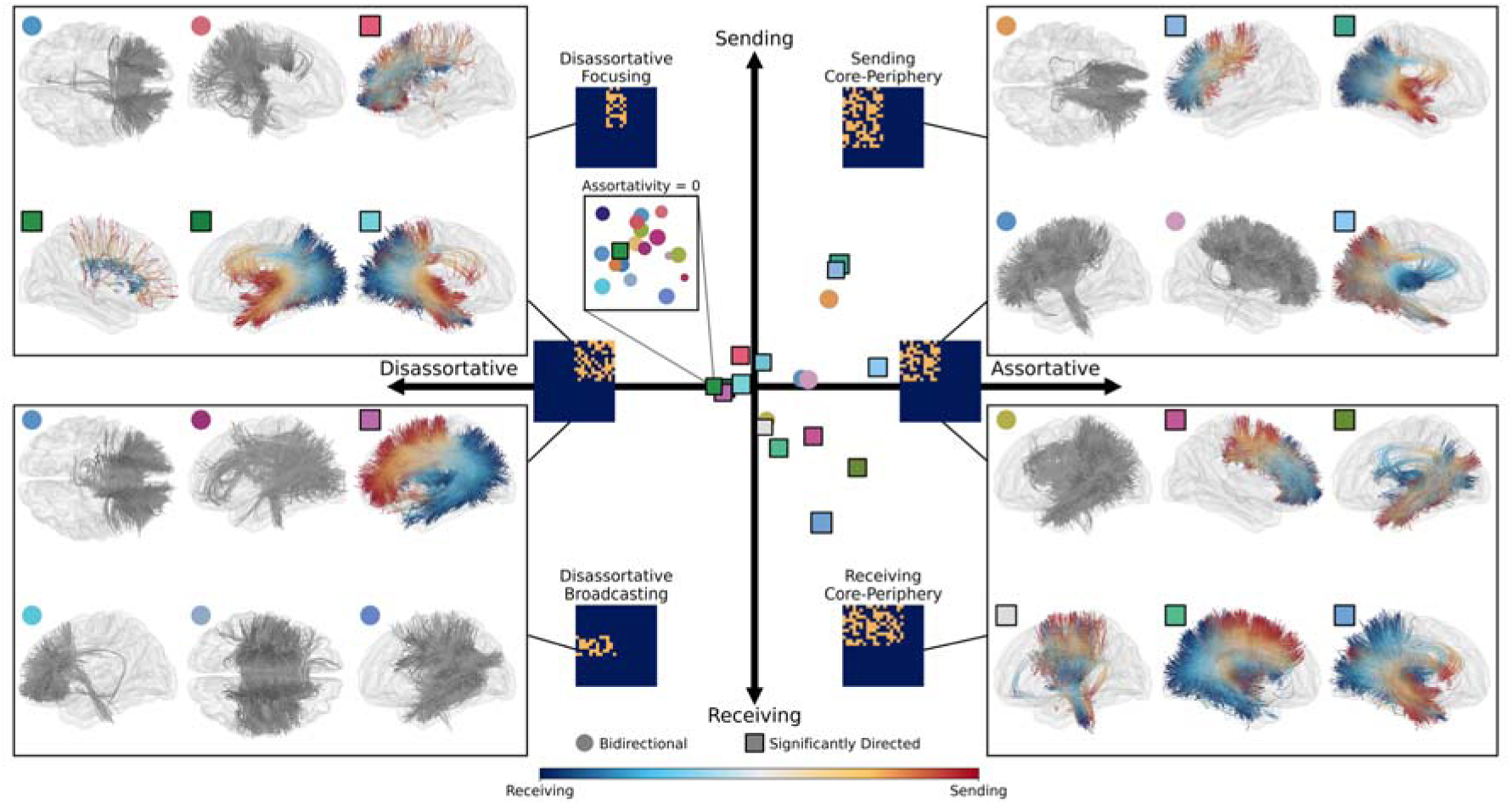
Assortative, Core-Periphery and Disassortative Bicommunities. The central scatter plot shows all *K*=35 bicommunities based on their assortativity (horizontal axis) and tendency to send or receive (vertical axis). Significantly directed and bidirectional bicommunities are shown with square and circle markers respectively. Marker colors reflect the colors of the clusters in Fig. 1. F and Fig. 2. Scatter size is proportional to the bimodularity index that captures the density of connections within a bicommunity. We show purely disassortative bicommunities in the inset box with title “Assortativity = 0”. Each quadrant is accompanied by a schematic binary adjacency matrix (0: dark blue, 1: yellow) that illustrates the type of connection as well as for purely assortative or disassortative communication. Finally, white matter streamlines centroids (5 per edge) of 6 selected bicommunities are shown in each of the 4 boxes in which they are ordered to reflect each quadrant of the scatter plot. Streamline representations have a colored marker indicating their correspondence to the central scatter. Five centroids of streamlines are shown for each edge in the bicommunity with a color going from red (sending) to blue (receiving). Streamlines of bicommunities with no preferred direction are colored in gray.

### Dissecting Association Fibers into Directed Bicommunities

We previously discussed that the structural wiring captured by the connectome edges can be viewed as sub-parts of white matter pathways. After validation of the directionality of bicommunities and fixing *K*=35, we now focus on these groups of edges and explore their neuro-anatomical and functional relevance. Utilizing the SCIL atlas of segmented white matter bundles (46,47), we compute the edge-to-bundle similarity for each of the graph edges defined as the inverse minimum average direct-flip (MDF) distance between streamlines (48). We then investigate how this similarity is reflected at the level of the bicommunities to identify the anatomical bundles most represented among its clustered edges (Fig. S7). Furthermore, we extract a network representation for each bicommunity by aggregating their connections based on the overlap between their sending (or receiving) nodes and 7 resting state functional networks (49). That way, each bicommunity can be expressed as a combination of anatomical bundles that describe directional communications between brain networks (Fig. 5, S4, S8). Bicommunities with preferred directions mostly describe parts of the association fibers, include parts of projection fibers (Fig. 5. G, H), and are coherently found in both hemispheres. We recognize well-defined combinations of the arcuate and superior longitudinal fasciculi, which sends from the frontoparietal and default mode back to the somatosensory and dorsal attention networks (Fig. 5. A, B). We further observe focused afferent connections towards the central visual cortex from temporal, dorsal and somatosensory areas along the inferior longitudinal fasciculus and the optical radiation (Fig. 5. C, D). Besides overall similarities between the two hemispheres, projection fibers with preferred direction are only found in the left hemisphere and capture both the afferent connections to the visual and dorsal networks and the efferent connections from the somatosensory area down towards the brainstem (Fig. 5. G, H). Finally, some other bicommunities describe similar pathways with opposite directions but do not show significant edge asymmetric (Fig. S4).

**Figure 5.**
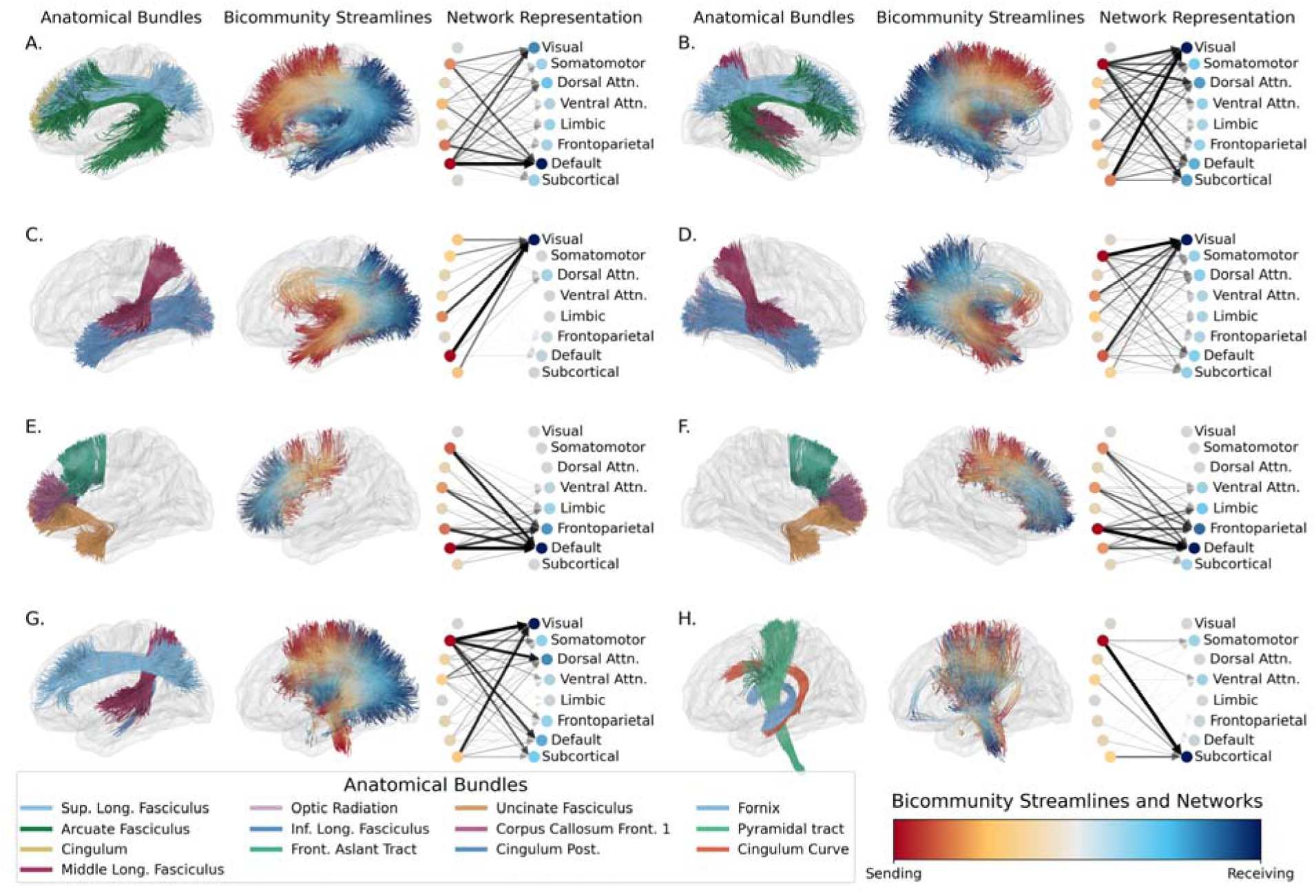
Bicommunities Capture Function Specific Anatomical Fiber Bundles. Anatomical bundle representation of 8 selected bicommunities (out of K=35) with significant asymmetry and high overlap with fiber bundles. For each of them (indicated with letters from A to H), we show: (**first column**) the most similar anatomical bundles; (**second column**) 5 streamline centroids for each edge in the bicommunity that are colored based on the which end of the streamline is sending (red) or receiving (blue); (**third column**) then sending (left, red) and receiving (right, blue) networks that are mapped by that bicommunity. For the network representation, edge width and opacity are proportional to the connectivity strength. Circles opacity represents the proportion of networks that belong in the sending or receiving nodes. Streamline and network directionality have been adapted to reflect the preferred direction (flipped if asymmetry < 0.5).

## Discussion

In this work, we revisit directed brain networks through the lens of bicommunities. Starting from a whole-brain structural connectome capturing the anatomical white matter scaffold, we add edge directionality from resting-state effective connectivity. Identifying bicommunities, that are meaningful groups of directed edges (42), we demonstrate that a majority of these pathways have significant edge asymmetry and are thus informed with a preferred direction that appears to be along the temporal-occipital-frontal axis. We simultaneously assess the relevance of connectome directionality and bicommunities, revealing that they meaningfully aggregate edge information consistent across independent estimations of directed communication and not detectable through edge-wise comparisons alone. We further confirm that brain communication patterns are not limited to densely connected components but rather follow a gradual spectrum from assortative to disassortative organization, with intermediate core-periphery structure. Finally, anatomical bundles of white matter (46,47) are well captured within specific bicommunities with the novelty of being informed with send-receive asymmetry, thus providing a rich perspective of how information may flow within and between functional networks.

### Data-Driven Bicommunities Capture Directed and Function Specific Pathways

By aggregating regional attributes and integrating edge directionality, bicommunities are holistic representations of network communication. While they are inherently informed about the white matter and functional organization of the human brain through the connectome, they naturally highlight coherent streams of information flow that map onto known segmented tracts (50). Overall, we observe an axis of preferential directionality that appears to support feedforward transmission of stimulus-driven sensory signals toward the association/executive cortex. Furthermore, bicommunities provide a basis for inferring potential functional specialization in groups of known white matter fiber bundles. Specifically, we find large overlap with the arcuate and superior longitudinal fasciculi in each hemisphere in bicommunities that describe broad communication from frontal to occipito-temporal and parietal areas (Fig. 5. A, B). These patterns map communication within and between default mode and attention networks, respectively, which could reflect the feedforward linguistic integration along the dorsal language pathway (51). On the other hand, core-periphery communication from temporal components of the DMN (medial temporal lobe) to occipital areas (visual cortex) through the inferior longitudinal and middle longitudinal fasciculi (Fig. 5. C, D) may represent crucial pathways for memory-guided visual processing, mental imagery, and episodic memory retrieval (52). In other bicommunities (Fig. 5. E, F), we observe short fibers bridging the somatosensory area to the frontoparietal network and further to the frontal cortex along the frontal aslant tract which could describe sensory-driven executive control. Finally, the last set of highlighted bicommunities, highlight two distinct directions of somatomotor-subcortical pathways that play key roles in sensory integration and movement coordination, supporting the notion that fibre directionality is shaped by functional demand. Differences in lateralization and network properties (sending and receiving core-periphery in left and right hemisphere respectively, Fig. 4), emerge in certain bicommunities (e.g., Fig. 5. E, F). This could reflect left/right asymmetry which is known in certain brain functions, such as, in this case, right hemisphere dominance for top-down executive control processes (53). While the discussed representations remain coarse (K=35), we demonstrate that bicommunities aggregate directional information to a scale that is biologically relevant and where modalities converge. Indeed, they provide an intuitive framework for identifying directed brain pathways, within which interpretations of functional specialization arise naturally.

### Coherent Models of Directed Brains

Integrating directionality in connectomes has shifted from a theoretical concept to more empirically grounded frameworks in which consistent patterns of edge asymmetry emerge across multiple modalities and timescales. By combining structural architecture with functional asymmetry, our approach aligns with this transition and shares similarities with existing models of directed brain connectivity (23,17,20,27). A general conclusion across DCM, Granger causality, or communication models is that neural architecture follows a particular hierarchy from sensory systems to integrative hubs (6). In detail, sensory areas are commonly identified as senders that broadcast information via feedforward loops to association regions (e.g., DMN), which are characterized by more incoming connections (17,20). This is reflected in some of the highlighted bicommunities in which edges point towards the visual cortex and are significantly stronger than their outgoing counterparts. We observe preferred directions from the temporal lobe passing through occipital areas and from somatosensory to frontal regions that are aligned with the sensory-association axis (6). Our results further indicate that large-scale white matter pathways with opposite directions may be passing through similar edges, hinting that communication efficiency within shared bundles depends on the direction and the associated functional demands. This is coherent with recent findings using whole-brain Granger causality during task fMRI showing that the flow of information from typically receiving regions is rebalanced depending on the task (54). In sum, similar organizational principles in directed brain graphs emerge from different methodological perspectives and modalities. Such a convergence hints that the observed and proposed directed asymmetries reflect biological constraints in the wiring and dynamics of the human brain.

### Mapping Brain Hierarchies Through Edge Communities

The detection of clusters of edges, or bicommunities, supports that brain community structure is not restricted to densely connected areas, but includes bipartite connectivity patterns (disassortative), in which parallel white matter pathways connect common origins and targets. This reflects organizational principles of white matter connections that span a continuum from functional assortativity (e.g., within sensory systems) to disassortativity (e.g., sensory-to-associative), which, in turn, may dictate the hierarchy of cognitive processes (10,39,55). Commonly found in human connectomes (56) and in other species (11,57), long-range disassortative connections are central to the efficient mapping between assortative hubs (58). Indeed, by linking low-degree sensory regions to highly connected association hubs, these bipartite-like and core-periphery structures constitute network-level evidence of the hierarchical information flow within the brain (13,56). Furthermore, white matter pathways are not isolated wires, but instead aggregate into dense bundles of fibers with shared sources, targets, and functional purposes (38,50). This is naturally captured by bicommunities in which connections are grouped if they share similarities in their sending and/or receiving nodes (42). We demonstrate that white matter pathways within bicommunities are not only biologically viable, but also function specific, with directional organization that may reflect the computational demands of the systems they mirror. This is supported by our results showing that the streamline representations of bicommunities, which we remind are purely data-driven, closely match known anatomical bundles and map directed information between specific functional networks. All in all, we establish that the presented bicommunities are biologically plausible aggregates that respect functional hierarchy and are mapped onto anatomical landmarks which could reflect both developmental constraints and optimizations for efficient information flow (59).

### Bridging Non-Invasive Connectomics with Invasive Validation

Structural and functional correspondence is not new when assessing the biological validity of models of directed connectivity (20,23,27). But alignment between brain-directed connectivity and anatomy alone is not sufficient to fully validate directed inferences and, as done in animal models, the gold standard stands in direct comparison between inferred directionality (as in EC) and electrical propagation (13,29) or circuit-level causality (30). This was, however, only achieved in clinical settings for humans, or only at a limited scale (14,60,61). Using invasive recordings of electrical propagation, we demonstrate that validation of the connectome directionality cannot be achieved at the level of single edges. This is not surprising as CCEPs are unevenly sampled and sparser than the structural connection (31). We, however, observe that a significant correspondence between the two modalities can be found when seen through the lens of bicommunities. This further hints that, with meaningful aggregates, patterns of directed connectivity are replicable across different dynamics and that there exists a scale at the crossroad where non-invasive and invasive measurements can be compared. It also demonstrates that the presented edge communities are not statistical artifacts or biased by methodological constraints but rather represent fundamental organizational units of directed wiring within the brain. This validation represents a key step toward human-level evidence that whole-brain directed connectivity inferred from non-invasive imaging aligns with invasive measures of electrical propagation.

The validation of bicommunities as a robust framework for dissecting directed brains opens several promising research directions. First, the integration of feedforward and feedback flow across cortical layers (e.g., with fMRI) could further unfold the hierarchical organization of connectomes by revealing which layer drives the asymmetric communication (62). Second, individualized connectomes suggest that population-averages architectures coexist with subject-specific connectivity profiles that reflect differences in anatomy and function. Therefore, investigating which bicommunities are shared across individuals, and which are not, could distinguish core directed architecture from individual variation and shed light on how development may have shaped directed connectomes. Finally, mapping how directed brain pathways may emerge during childhood or reorganize following injury appear as compelling developmental and clinical applications. These extensions thus situate bicommunities as a pivotal framework for understanding brain development, adaptation and disease.

We list a few methodological limitations to the presented work. First, effective connectivity (estimated from fMRI) operates at different timescales (seconds against milliseconds) and characterizes different communication processes (lagged synchronization against electrical propagation) than invasive recordings. However, by comparing asymmetry rather than connectivity strength (or delay), our approach is scale invariant and captures potential imbalances between outgoing and incoming edges. Furthermore, evidence supports that fMRI hemodynamic timing accurately reflects neural activation sequences despite different timescales, hinting that asymmetries are thus comparable (63). Biologically, these asymmetries likely reflect underlying developmental optimization through myelination of specific white matter pathways, which, in terms, may exhibit imbalance in efficiency that is similarly observable across different temporal scales (64). Second, we ensure robustness of edge clusters through consensus and hierarchical clustering, which serve as methodological validation of parameters (e.g., number of clusters *K*). However, the bicommunities remain data-driven, and edges could still be mislabeled. Nevertheless, we demonstrate that the highlighted white matter pathways are neurobiologically relevant and robust across granularity. Third, the structural backbone of our directed connectome is computed from diffusion MRI and tractography, which are known to have low specificity (65). However, we only consider a binary structural graph after conservative thresholding of both the average number of connecting streamlines and participants with existing connections (45). We further note that projection fibers are underrepresented within the connectome which could explain that few bicommunities capture such pathways. Finally, each of the three discussed modalities are aggregated from different populations. Specifically, the invasive CCEPs are measured in patients with epilepsy as part of the F-Tract consortium (31), while structural connectivity is computed from a healthy population. Similarly, while statistical significance is already implemented in the peak detection procedure within F-Tract, we thresholded out connections with few measurements or implants to only retain reliable estimates.

To conclude, we presented a whole-brain directed connectome in which asymmetric white matter pathways are informed by effective connectivity. We demonstrated that bicommunities naturally aggregate edges into large-scale pathways according to their send-receive asymmetry profile and reveal hierarchical information flow that is concealed by traditional nodal methods. Validation against intracranial recordings supports the bicommunity structure and connectome directionality. Finally, known anatomical bundles of white matter are identified and, for some, informed with a preferred direction which reflects their overall directed mapping across functional networks. All together, these advances position directed connectomics and bicommunities as a new tool for understanding brain organization.

## Materials and Methods

### Merging Effective and Structural Connectivity in a Directed Connectome

The structural part of the directed connectome comes from a population-level atlas of white matter streamlines from diffusion MRI (45). Specifically, 129 (114 cortical, 7 bilateral subcortical regions, and the brainstem) nodes from the Lausanne 2018 parcellation (scale 2) (66), which are connected by an edge if there are at least 5 streamlines present connecting the region pairs in at least half of the subjects (i.e., in more than 33 participants). Corresponding streamlines are then aggregated, for each connection, into centroids for visualization purposes. To reduce the effect of bidirectional connectivity strength, we estimate the connectome directionality as the edge-wise asymmetry level from directed whole-brain resting-state effective connectivity from regression Dynamical Causal Modelling (23,35). In detail, and for the remainder of the study, we define asymmetry of an edge e*_ij_* between nodes *i* and *j* as the ratio 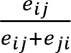 that captures the proportion of outgoing strength. We note that only 4% of the edges had negative connectivity weights and were thus removed to ensure consistency with the asymmetry ratio. Furthermore, the original effective connectivity data were in the Schaefer 400 atlas with 7 additional bilateral subcortical regions (67). We thus mapped the original effective connectivity matrix to our current parcellation by computing a network correspondence matrix based on the proportional overlap between Schaefer 400 and Lausanne 2018 parcels. Then, we build the connectome where edges can only exist if there is a white matter connection (binary structure), and with weights given by the asymmetry proportion estimated from effective connectivity.

### Hierarchical Detection of Directed-Graph Communities

We leverage the recent bimodularity (42) framework to extract directed brain communities defined as clusters of graph edges. In short, edges are projected onto an embedding space that captures their sending and receiving patterns defined by the singular value decomposition of the directed modularity matrix (68). In such a space, edges are grouped using K-means to represent directed communities named bicommunities. We ensure stability of the clusters through the means of consensus clustering by repeating initializations of K-means for varying numbers of clusters (*K*=10 to *K*=80, 50 initializations each). Furthermore, stable numbers of clusters are highlighted with hierarchical clustering applied to the consensus matrix, which measures the likelihood for two edges to fall in similar clusters.

### Non-Parametric Testing for Significant Asymmetries

To assess the statistical significance of directionality, we first aggregate the edge weights within a bicommunity as the median edge asymmetry. Then, we build a null distribution of these asymmetries by randomly swapping the weights of half of the connectome edges with their corresponding edge going in the opposite direction (N_Permutations_=10000). This approach essentially identifies bicommunities with statistically significant asymmetry levels that reject the null hypothesis of “edge directionality does not matter”. P-values are computed as the ratio between the number of random asymmetries higher than the true one and the total number of random permutations, with an added normalization factor of 1. Finally, p-values are corrected for multiple testing using the Bonferroni correction with a number of tests equivalent to the number of bicommunities (*K*=35 in Fig. 2) and considering a significance threshold of α=0.05.

### Assortativity Level and Send-Receive Dominance

We estimate the assortativity of a bicommunity as the similarity between its sending and receiving nodes. In detail and for each cluster, sending and receiving vectors are built as the proportion of outgoing (or incoming) edges that belong in the cluster and represent a probability for an edge to be a sender (receiver) in the bicommunity. Then, we estimate the assortative connectivity pattern that would be given by connection within the sending nodes or within the receiving ones as the outer product of each vector with themselves. We define the assortativity level as the average overlap between the bicommunity edges and the ideal sending and receiving assortative patterns. Therefore, if the proportional overlap is high in both cases, assortativity will be high. However, if one overlap is higher than the other it means the bicommunity edges connect most nodes of either the sending or receiving set, but not the other. In other words, these connections appear as core-periphery types of connectivity and map a higher number of sending nodes to few receiving ones or the opposite. We thus quantify this imbalance as the difference between the edge overlap with sending and receiving assortative patterns. Positive values indicate more sending than receiving nodes and capture focusing patterns of communication; bicommunities with negative values thus describe broadcasting edges. Finally, if small or no overlap is found for both sending and receiving patterns, the bicommunity is considered as disassortative with assortativity being close to zero.

### CCEPs from the F-Tract Consortium

For validation, we included directed cortico-cortical evoked potentials (CCEPs) from stereo-encephalographic (SEEG) responses to a direct electrical stimulation (12,31). The precise procedure for temporal peak detection and estimation of electrical signal propagation is outlined in earlier work on the F-Tract dataset (https://f-tract.eu) (31). Briefly, SEEG recordings in patients with drug-resistant epilepsy that were awake at rest were aggregated from data from 21 centers. The SEEG data analysis was as follows: first, bad channels were identified with a machine learning algorithm supervised by trained personnel. Then, according to standard procedure, the signals were referenced to the bipolar montage to minimize volume conduction effects. Third, short (4 ms) stimulation artefacts were removed by local signal blanking and interpolation. Then, band pass filtering was applied between 1 and 45 Hz and responses to all stimulation pulses within one run were averaged, baseline corrected and z-scored with respect to [−200, −10] ms time before stimulation. Such averaged response was considered significant if its absolute value exceeded a z-score threshold Z=5 within a [0, 50] ms time window to avoid the contribution of indirect communication (axonal propagation through multiple regions). They were obtained for 10551456 recordings coming from 90653 stimulations with mean current intensity 3.4 mA +/- 1.3 std, mean pulse width 1 ms +/- 0.5 std and frequency 1 Hz. In 75% (25%) of cases stimulations were performed in the biphasic (monophasic) manner. In total, they correspond to 1307 implantations (641 males, 660 females, 6 unknown) performed in 1236 patients: 602 male (mean age 25 years +/- 14 std), 628 female (mean age 26 years +/- 13 std), 6 unknown (mean age 30 years +/- 10). Among all features in the F-Tract database, we included temporal features such as the onset delay (time between stimulation and the passing of the statistical threshold), peak delay (time between stimulation and the first peak of the response), as well as latency start (time between the stimulation and the last local minimum before the peak). Also, we considered non-temporal features such as the response amplitude (height of the first peak). These measurements were then aggregated for each connection between pairs of regions in the atlas of our connectome to build F-Tract matrices of connectivity for each feature. Finally, to further reduce noisy recordings, we only considered connections in which at least 50 recordings were present from at least 3 unique implantations (69).

### Validating Connectome Asymmetry with the Invasive F-Tract Dataset

We first compute edge asymmetry levels for each F-Tract connectivity matrix in which high asymmetry either describes longer delays (temporal features) or higher amplitude in the response. To compare these asymmetry levels with our directed connectome, we use the Spearman correlation at the level of the graph edges, graph bicommunities, and when shuffling the cluster assignment of graph edges. This shuffling thus creates a null distribution of correlations between the connectome and CCEPs edge asymmetries in settings in which shuffled bicommunities are clusters of random graph edges (N_Permutations_=4999). Then, we compare the correlations obtained with true bicommunities with the random ones and evaluate their statistical significance. P-values are computed with the similar approach as for the significance of bicommunity edge asymmetry. Finally, p-values are corrected for multiple comparisons using the false discovery rate (FDR) correction (70).

### Anatomical Bundle and Functional Network Representation of Bicommunities

We evaluate how segmented white matter bundles from the SCIL atlas (46,47) are represented within bicommunities. First, we estimate the similarity between connectome edges and anatomical bundles using minimum average direct-flip (MDF) distance which captures the minimum spatial distance between groups of streamlines (48). Second, we build an edge-to-bundle matrix of similarity in which entry A*_m,n_* is the inverse MDF distance between the streamlines of edge *m* and atlas bundle *n*. Finally, by aggregating entries of the similarity matrix for all edges within a bicommunity, we obtain a bicommunity-to-bundle similarity matrix. We show the edge-to-bundle and bicommunity-to-bundle similarities for each bundle of the atlas in Fig. S8. Similarly, we compute a matrix of proportional overlap between regions of our parcellation and the 7 Yeo resting-state networks (49). To do so, we compute, for each atlas region, the proportion of voxels that overlap with functional networks which is then summarized in a region-to-network matrix. From this, we build a network definition of the sending and receiving nodes of a bicommunity which summarizes the networks from which these edges originate and to which they project.

## Funding

This work was supported in part by the Swiss National Science Foundation through the Sinergia project “Precision mapping of electrical brain network dynamics with application to epilepsy,” grant No. 209470, and the regular project with grant No. 207493.

## Author contributions

Conceptualization: AC, CHMC, FS, MGP, DVDV

Methodology: AC, MJ, OD, MGP, DVDV

Investigation: AC, FS, MJ

Visualization: AC, FS, YAG

Supervision: MGP, DVDV

Writing—original draft: AC

Writing—review & editing: AC, CHMC, FS, MJ, YAG, SA, AS, OD, IJ, PH, MGP, DVDV

## Competing interests

The authors declare that they have no competing interests.

## Data and materials availability

Previously published data were used for this work (31,35,46,47). In detail, the probabilistic atlas of white matter connection can be accessed from https://github.com/connectomicslab/probconnatlas (45). Estimates of effective connectivity are found in the repository at: https://zenodo.org/records/14034721 (35). Invasive CCEPs from the F-Tract consortium are available on the EBRAINS platform https://search.kg.ebrains.eu/instances/41db823e-7e1b-44c7-9c69-eaa26e226384 (31). Finally, segmented white matter fiber bundles are accessible from the RecobundleX pipeline repository https://github.com/scilus/rbx_flow (46,47). All code and data are openly available at: https://github.com/MIPLabCH/Bimodularity. All data needed to evaluate the conclusions in the paper are present in the paper and/or the Supplementary Materials.

## Supplementary Materials

Available in a separate file.

## Notes

### Competing Interest Statement

The authors have declared no competing interest.

