## Supplementary Material for "Directed Brain Connectomics Revealed by Bicommunity Structure"

Alexandre Cionca *et al.*

**This PDF file includes:**

Supplementary Text

Figs. S1 to S8

Supplementary Text

Abbreviations of the atlas of anatomical bundles of white matter

Below, we summarize the abbreviations of the SCIL (46,47) atlas of white matter bundles which might be used in the main text (as reported in (47)):

AC: Anterior commissure;

AF: Arcuate fasciculus;

CC_Fr_1: Corpus callosum, Frontal lobe (most anterior part);

CC_Fr_2: Corpus callosum, Frontal lobe (most posterior part);

CC_Oc: Corpus callosum, Occipital lobe;

CC_Pa: Corpus callosum, Parietal lobe;

CC_Pr_Po: Corpus callosum, Pre/Post central gyri;

CC_Te: Corpus callosum, Temporal lobe;

CG: Cingulum;

FAT: Frontal aslant tract;

FPT: Fronto-pontine tract;

FX: Fornix;

ICP: Inferior cerebellar peduncle;

IFOF: Inferior fronto-occipital fasciculus;

ILF: Inferior longitudinal fasciculus;

MCP: Middle cerebellar peduncle;

MdLF: Middle longitudinal fascicle;

OR_ML: Optic radiation and Meyer's loop;

PC: Posterior commissure;

POPT: parieto-occipito pontine tract;

PYT: Pyramidal tract;

SCP: Superior cerebellar peduncle;

SLF: Superior longitudinal fasciculus;

UF: Uncinate fasciculus.

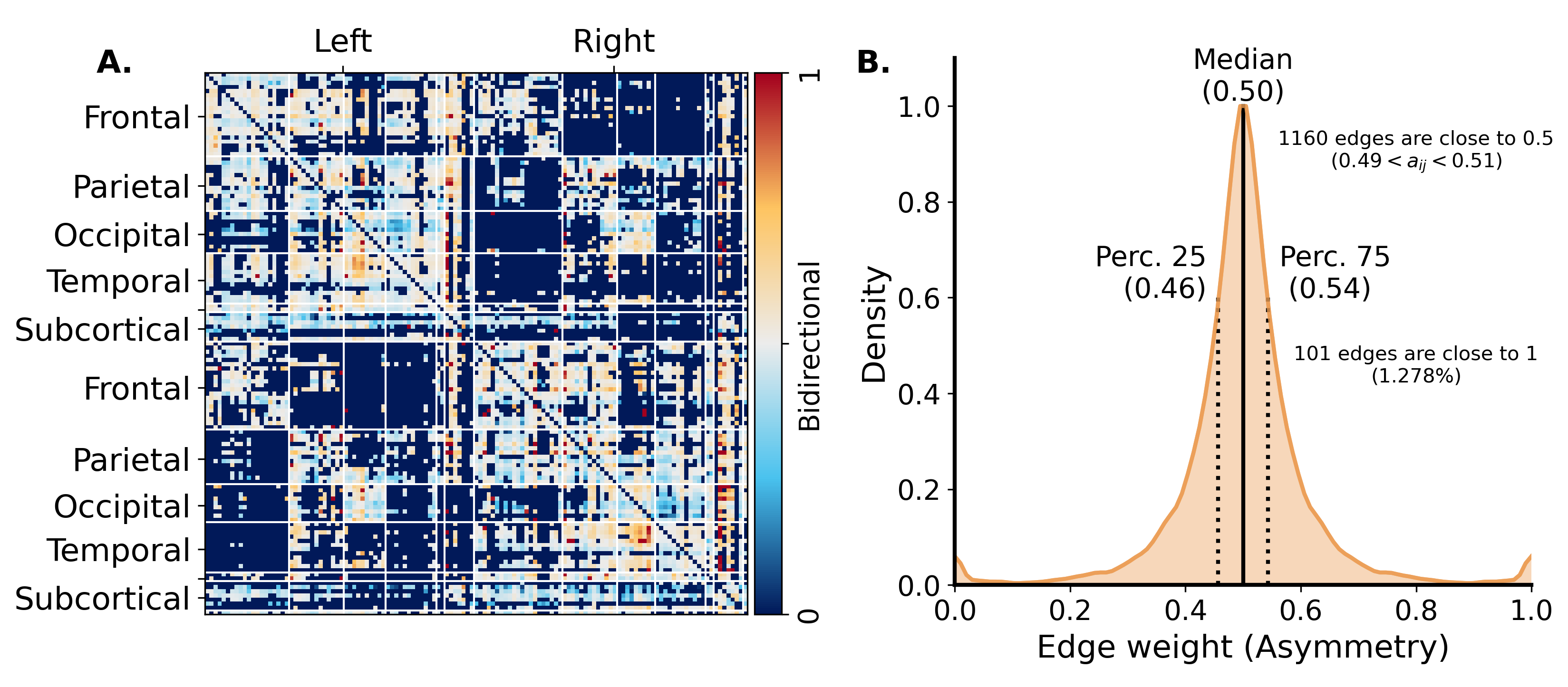

Fig. S1. Summary of the presented directed connectome.

**A**) Adjacency matrix of directed connectivity with nodes ordered by lobe and hemispheres. Connectivity value of 0.5 is equivalent to bidirectional connections. **B**) Distribution of edge weights in the connectome. One half of the connections are within the [0.46, 0.54] range which tend to be bidirectional while the other half appear as asymmetric. On one hand, 1160 edges (out of the 7906, 14.67%) have directionality close to 0, indicating pure bidirectional connections. On the other hand, 1.278% of connection are purely directed (edge weight close to 1).

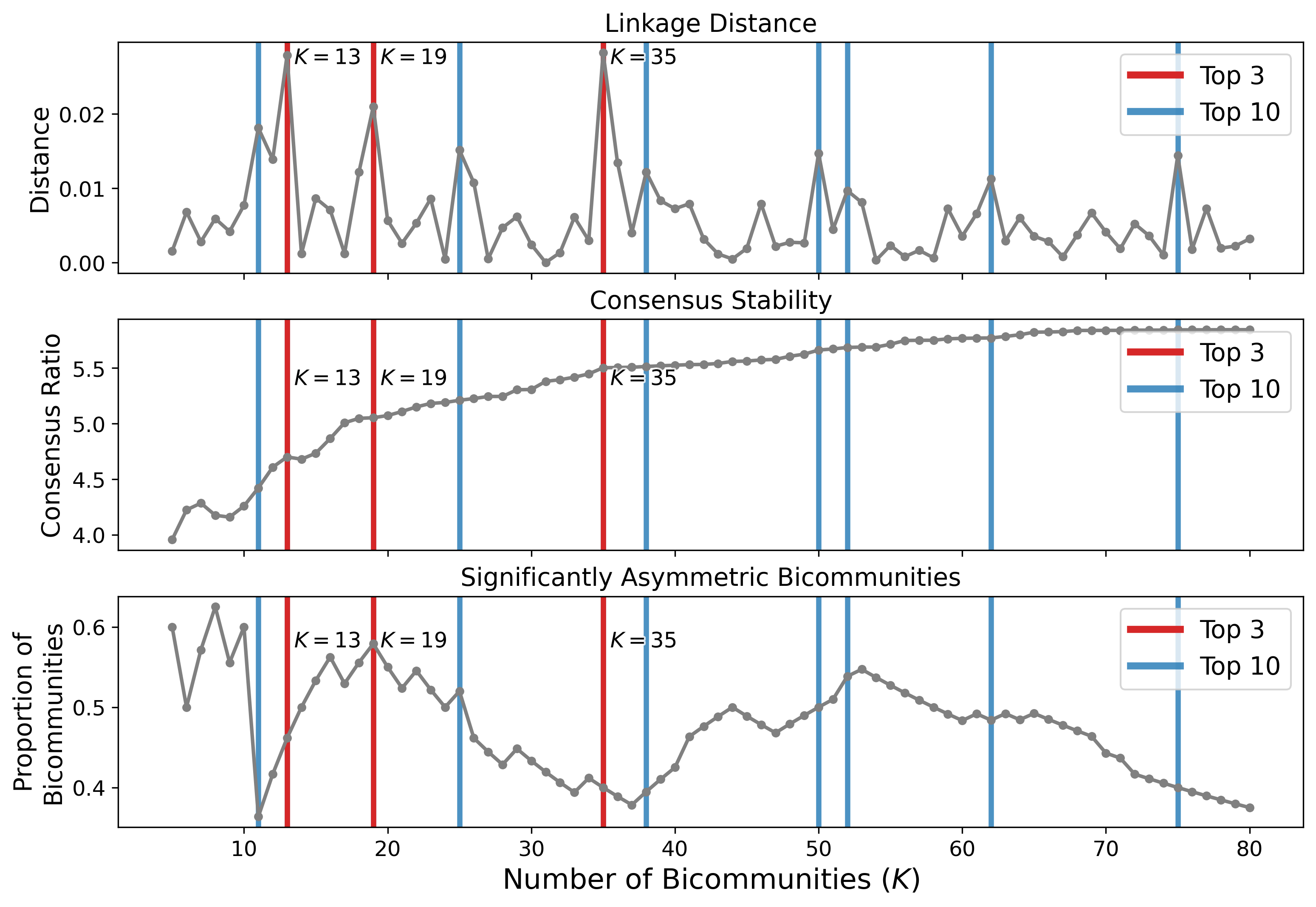

Fig. S2. Stability of hierarchical consensus clustering and significance of asymmetry

Stability of edge clustering and proportion of asymmetric bicommunities for a range of number of edge clusters (*K=5* to *K=80*). For all top 10 (blue) and top 3 (red) most stable number of bicommunities, we show stability criteria given by the hierarchical linkage distance (top) and the ratio between within and between cluster co-clustering probability (value of the consensus matrix, center). Finally, we show that the proportion of statistically significant bicommunities (as estimated through permutation testing, N permutation = 10000) remain within a similar range for stable clustering.

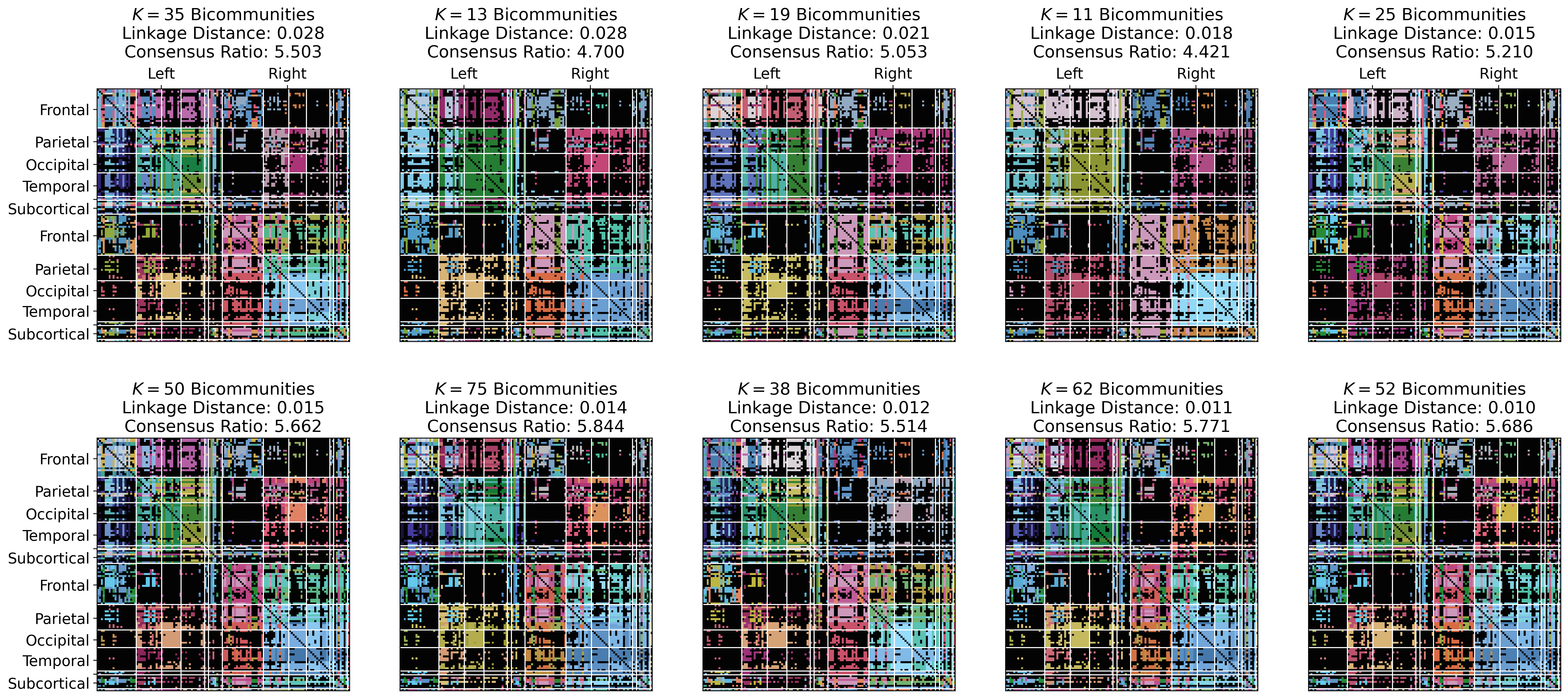

Fig. S3.

Cluster assignment of graph edges for each of the top 10 most stable number of bicommunities (as estimated through hierarchical clustering). The clustering is summarized in matrices in which row and columns represent the sending and receiving regions respectively which are similarly ordered by hemisphere (left then right) and lobe.

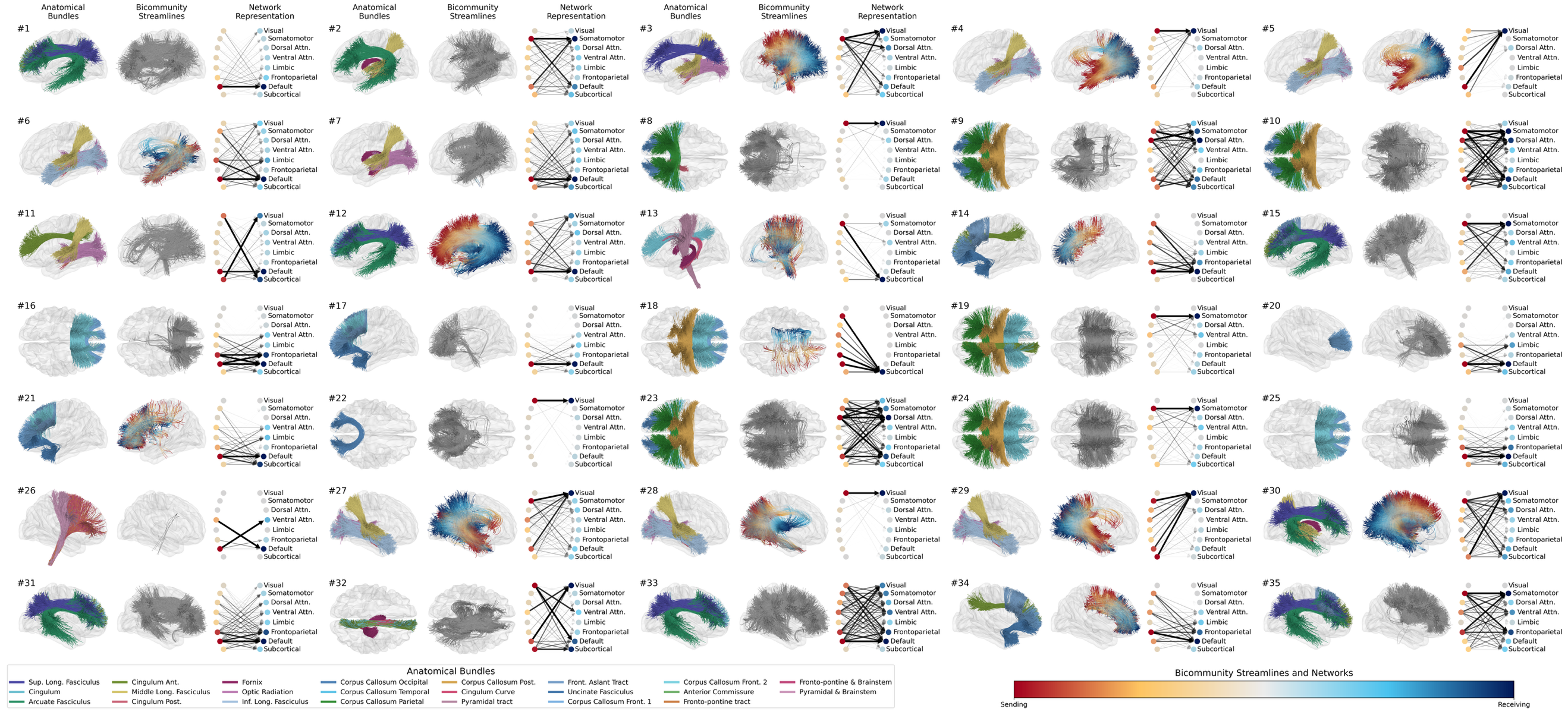

Fig. S4. Fiber bundles and resting state network representation of all *K*=35 bicommunities.

For each bicommunity indicated with numbers from 1 to 35, we show: (**first column**) the most similar anatomical bundles; (**second column**) 5 streamline centroids for each edge in the bicommunity that are colored based on the which end of the streamline is sending (red) or receiving (blue). Gray streamlines indicate bicommunities with non-significant edge asymmetry; (**third column**) then sending (left, red) and receiving (right, blue) networks that are mapped by that bicommunity. For the network representation, edge width and opacity are proportional to the connectivity strength. Circles opacity represents the proportion of networks that belong in the sending or receiving nodes. Note that streamline and network directionality have been adapted to reflect the preferred direction (flipped if asymmetry < 0.5) and that edges are reciprocal (and thus undirected) for bicommunities that do not show significantly asymmetric edge weights.

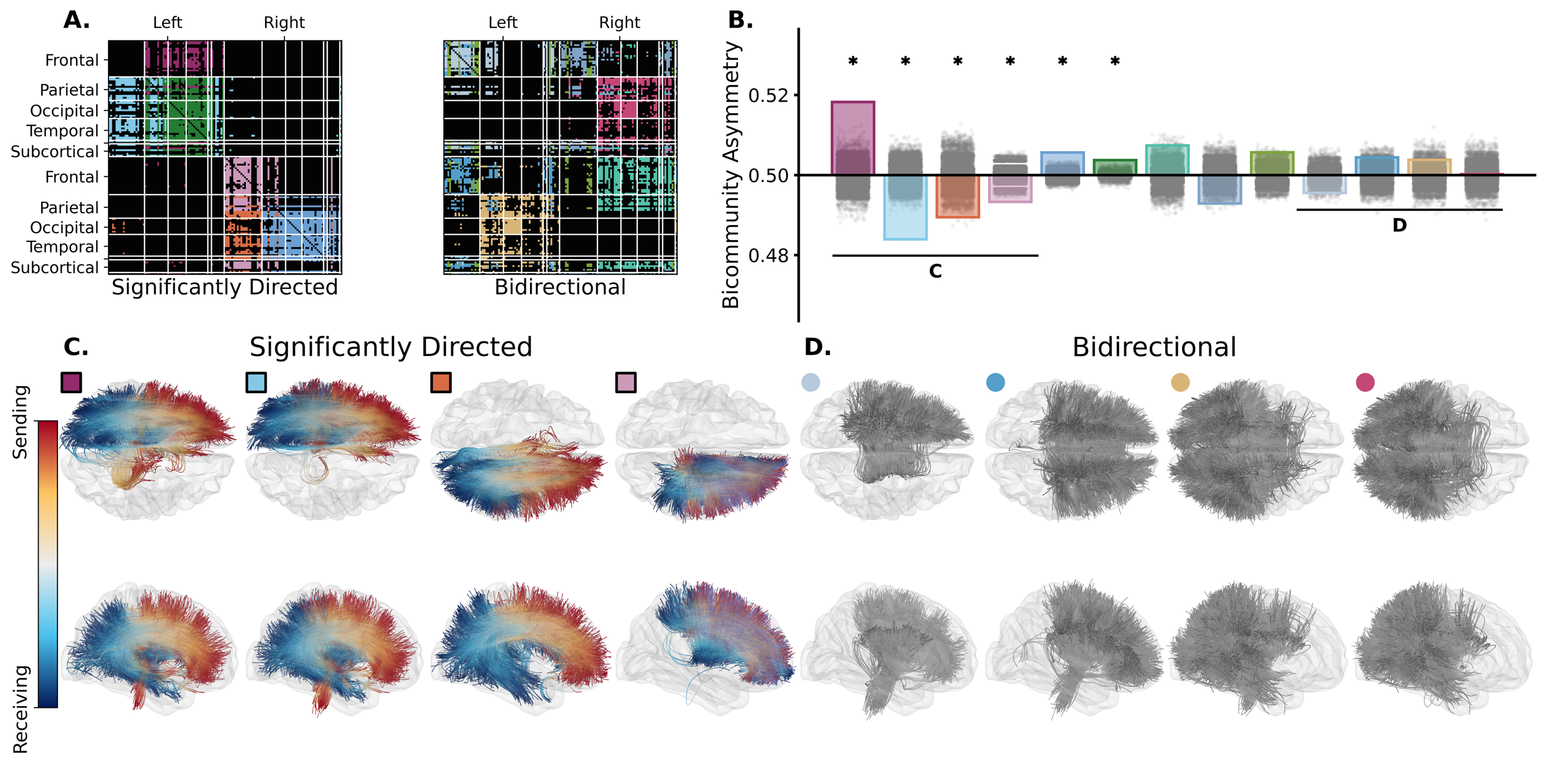

Fig. S5. Significant edge asymmetry for *K*=13 bicommunities.

**A**) Edge cluster matrix showing the cluster assignment of edges to individual bicommunities separated in significantly directed (left) or non-significant (right) edge asymmetry. Graph nodes are ordered by brain lobes (rows) and hemisphere (columns). **B**) Bar plots show the median edge asymmetry within each of the *K*=13 bicommunities with colors to indicate edge to cluster assignment as in **A**. Gray scatter points show the null distribution of asymmetry after reshuffling edge weight directions, and dark asterisks show significantly directed bicommunities (p-value < 0.05 after Bonferroni correction). **C-D**) White matter streamline centroids are shown for the 8 bicommunities with the highest (**C**) and lowest (**D**) median directionality (as highlighted with two horizontal lines in **B**) in the transverse (first and third rows) and sagittal (second and fourth rows). Colored markers show the bicommunity correspondence with **A** and **B** where squared and circle markers indicate significant and non-significant edge asymmetry respectively. Streamlines are colored based on which end is sending (red) and receiving (blue), or fully in gray for bidirectional bicommunities. Streamline directions have been adapted to reflect the preferred direction (flipped if edge asymmetry is < 0.5).
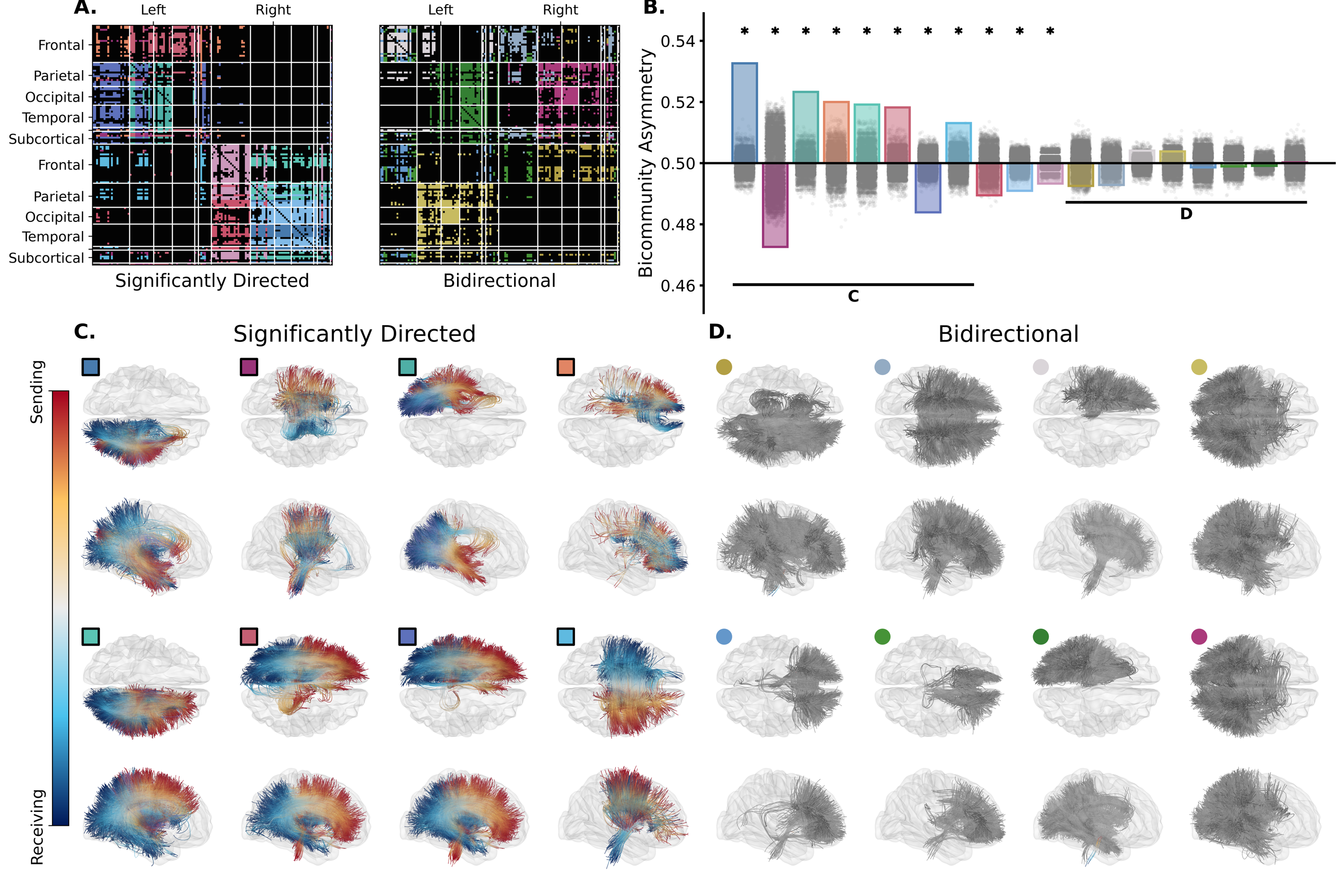

Fig. S6. Significant edge asymmetry for *K*=19 bicommunities.

**A**) Edge cluster matrix showing the cluster assignment of edges to individual bicommunities separated in significantly directed (left) or non-significant (right) edge asymmetry. Graph nodes are ordered by brain lobes (rows) and hemisphere (columns). **B**) Bar plots show the median edge asymmetry within each of the K=19 bicommunities with colors to indicate edge to cluster assignment as in **A**. Gray scatter points show the null distribution of asymmetry after reshuffling edge weight directions, and dark asterisks show significantly directed bicommunities (p-value < 0.05 after Bonferroni correction). **C-D**) White matter streamline centroids are shown for the 8 bicommunities with the highest (**C**) and lowest (**D**) median directionality (as highlighted with two horizontal lines in **B**) in the transverse (first and third rows) and sagittal (second and fourth rows). Colored markers show the bicommunity correspondence with **A** and **B** where squared and circle markers indicate significant and non-significant edge asymmetry respectively. Streamlines are colored based on which end is sending (red) and receiving (blue), or fully in gray for bidirectional bicommunities. Streamline directions have been adapted to reflect the preferred direction (flipped if edge asymmetry is < 0.5).
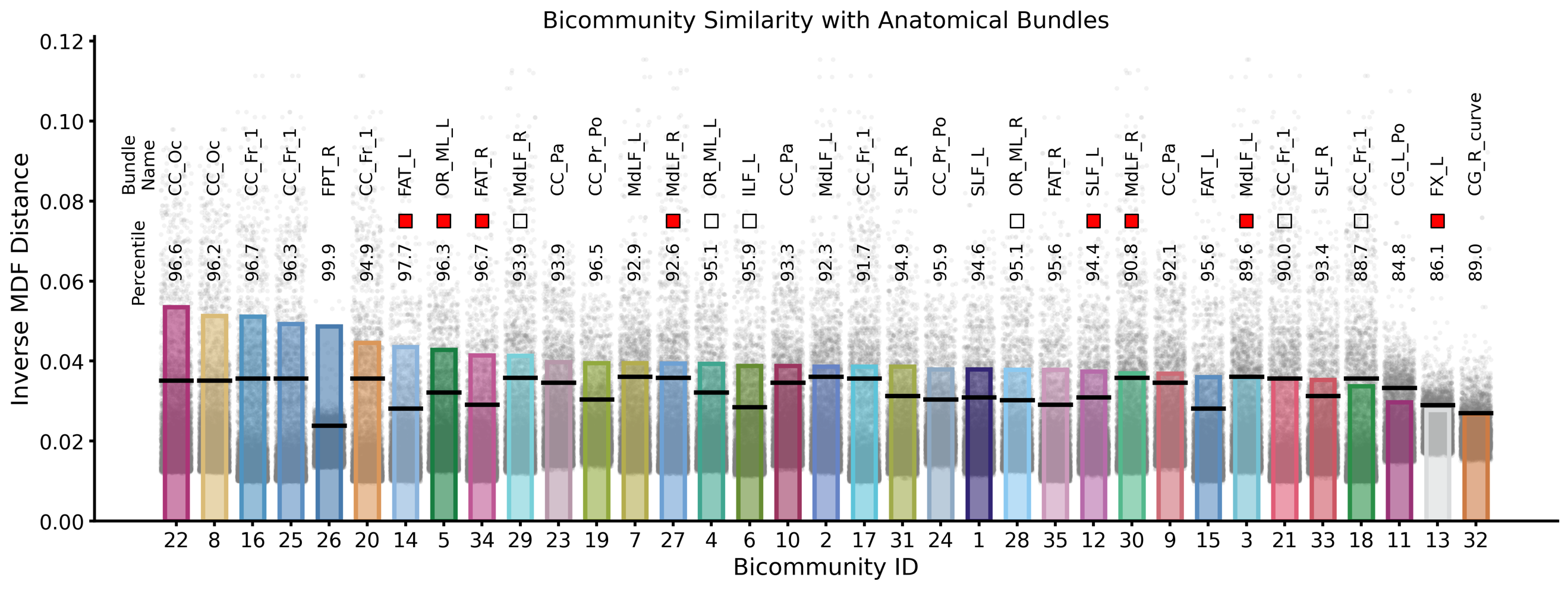

Fig. S7. Similarities between bicommunity streamlines and anatomical fiber bundles.

Inverse minimum average direct-flip (MDF)(48) distance between streamlines of bicommunities (5 centroids per edge) and segmented fiber bundles (50). For each bicommunity, we show the similarity with anatomical bundle that shows the highest inverse MDF distance (colored bar) with bundle label indicated on the top (“Bundle Name” legend). We show the inverse MDF distance between the selected bundle and steeamlines of each edge in the connectome with the gray scatter. The black horizontal bars mark the 90^th^ percentile of edge-to-bundle similarity and the numbers on the right of the “Percentile” legend indicates the percentile of the bicommunity-to-bundle similarity when compared to the overall distribution of edge-to-bundle inverse MDF distance for that specific fiber bundle. Finally, square markers indicate bicommunities with significantly asymmetric edge weights and are colored in red for those selected in Fig. 5 of the main text.

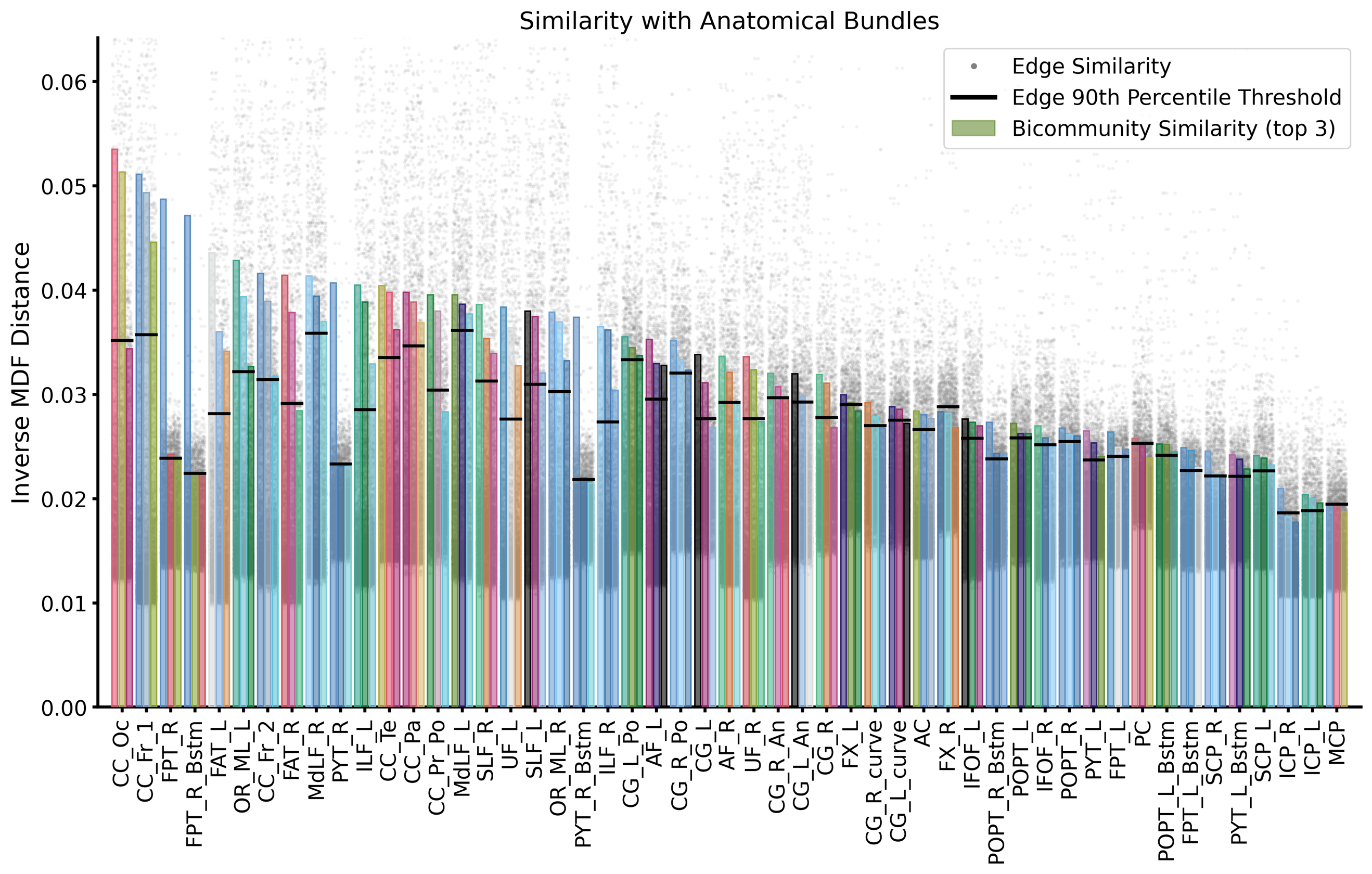

Fig. S8. Representativity of anatomical fiber bundles.

Similarities between connectome edges and segmented anatomical bundle of white matter fibers as captured by the inverse minimum average direct-flip (MDF) distance (48). We further overlay that same similarity, aggregated within each bicommunity, for the top 3 bicommunities with highest overlap. In detail, scatter dots represent the inverse MDF distance between the centroid streamline of each graph edge and anatomical bundle (as indicated in the horizontal axis). Each colored bar shows the same metric but averaged at the level of a bicommunity. Horizontal bars show the graph edge at the 90th percentile of similarity with anatomical bundle. We observe that, for most bundle, some bicommunities have higher similarity than the 90^th^ percentile threshold.
